# Individual Patient Data Analysis of Liver Lesion Dynamics: A Pan Cancer Analysis

**DOI:** 10.1101/2024.10.27.620503

**Authors:** Jake Dickinson, Heinrich J. Huber, Hitesh B. Mistry

## Abstract

There are now several databases available that enable the exploration of individual patient data from past Oncology clinical trials. These databases provide researchers with the opportunity to investigate the dynamics of individual lesions across various tumour sites within a patient and compare them across different patients. In this report, we specifically focus on liver metastases and examine the dynamics of individual lesions across different treatment and tumour types. Our findings reveal that when a lesion progresses early on, the rate of growth is significantly higher for liver lesions compared to other types of lesions, regardless of the treatment modality (chemotherapy, targeted therapy, or combination of both). These results indicate the need for further basic research on tumours and their microenvironment specifically within liver metastases. This analysis demonstrates how analysing open trial data can uncover new research prospects at the fundamental science level, thereby bridging the gap between laboratory research and clinical applications.

## Introduction

In the past, there were limited opportunities to explore individual patient data from historical phase III clinical trials, as this data was typically kept within companies and regulatory agencies. However, in recent years, there has been a rise in databases that provide researchers with access to such data (1– 3). This has resulted in interesting insights and articles that contribute to our understanding of diseases from a clinical perspective (4–7). In this report, our focus will be on the imaging data obtained from these databases, which comes from a series of studies across different tumour types. We aim to examine the kinetics of different lesions, with a specific emphasis on liver metastases, which is a significant prognostic factor in many indications (8–15). The rationale behind this research is to investigate whether a key metastatic site exhibits different kinetics compared to other lesions across various cancer indications and treatment modalities.

## Method

### Data

Data from individual patient imaging was obtained from ProjectDataSphere (PDS), encompassing five tumour sites: metastatic colorectal cancer (mCRC) (16), gastroesophageal junction (GEJ) (17), hepatocellular carcinoma (HCC) (18), non-small cell lung cancer (NSCLC) (19), and pancreatic cancer (PC) (20). Detailed information about each study can be found in the original trial publication cited, furthermore the PDS identifiers and hyperlinks for each study were [mCRC](https://data.projectdatasphere.org/projectdatasphere/html/content/309):309, [GEJ](https://data.projectdatasphere.org/projectdatasphere/html/content/130):130, [HCC](https://data.projectdatasphere.org/projectdatasphere/html/content/124):124, [NSCLC (Docetaxel)](https://data.projectdatasphere.org/projectdatasphere/html/content/133):133, [NSCLC (erlotinib)](https://data.projectdatasphere.org/projectdatasphere/html/content/115):115 and [PC](https://data.projectdatasphere.org/projectdatasphere/html/content/442):442. These particular studies were selected because they included imaging data recorded using RECIST (Response Evaluation Criteria In Solid Tumours) and provided specific anatomical location information. We specifically extracted longitudinal data on “Target Lesions” as defined by RECIST, which are the only lesion types recorded quantitatively over time. The data includes the longest diameter measurement of each lesion over time, along with its corresponding anatomical location

### Tumour metrics

The extracted tumour metrics include the initial lesion size measured in millimetres (mm) and the initial net rate, which is defined as the difference in lesion size between the pre-treatment visit and the first on-treatment visit divided by the time interval between the two visits. Within the datasets, lesions are categorized into three groups based on RECIST criteria during the initial imaging visit: PD (Progressive Disease), SD (Stable Disease), or PR/CR (Partial or Complete Response). This categorization is utilized to stratify the net rates of the lesions. Furthermore, the net rates are further stratified based on the anatomical location, specifically whether the lesion is located in the liver or elsewhere.

## Results

In the following sections, we examine the tumor lesion metrics for each specific cancer type and treatment, as outlined in the methods. It is crucial to recognize that the data used in these analyses were derived from randomized control trials; however, not all studies provided data for both arms.

### mCRC

Among the available data from various studies, the mCRC study stood out as the only one with data from two treatment arms: FOLFOX and FOLFOX + Panitumumab (EGFR antibody). This study also included information on the KRAS status of patients, which was investigated as a potential marker for resistance to the effects of Panitumumab. Figure 1 illustrates the changes in the average lesion size (as individual lesion identifiers were not available) from pre-treatment to the first on-treatment time-point (approximately 8 weeks), categorized by KRAS status. Figure 2 provides a breakdown based on lesion type, specifically liver versus other sites. To quantify the data presented in Figures 1 and 2, we refer to Tables 1 and 2. Table 1 demonstrates that the initial lesion size and kinetics, stratified by response type, are similar between the two treatment arms. However, when considering the KRAS status (as shown in Table 2), we observe that patients with progressive disease (PD) exhibit a higher net-rate in the FOLFOX + Panitumumab arm compared to the FOLFOX alone arm, specifically in cases of KRAS mutant versus wild-type. This indicates a clear interaction between KRAS mutational type and treatment response, with patients who do not achieve disease control (defined as stable disease (SD), partial response (PR), or complete response (CR)) experiencing faster progression when receiving Panitumumab. Further stratification by lesion location, specifically liver versus other sites (as demonstrated in Table 3 and Table 4), reveals that the disparities in PD net-rates between the two treatment arms are primarily observed in liver lesions.

**Table 1.** Calculated summary statistics (median, minimum and maximum) by treatment arm in mCRC patients.

| Variable | Class | FOLFOX alone | Panitumumab + FOLFOX | p-value |
| --- | --- | --- | --- | --- |
| Starting lesion size (mm) | - | 40 (20, 190) | 39 (20, 191) | - |
| Net rate (mm/day) | Overall | -0.121 (-0.898, 0.299) | -0.154 (-1.04, 0.323) | - |
|  | PD | 0.19 (0.0884, 0.299) | 0.208 (0.102, 0.323) | 0.417 |
|  | SD | -0.0819 (-0.435, 0.241) | -0.079 (-0.368, 0.0585) | 0.477 |
|  | PR/CR | -0.217 (-0.898, -0.0848) | -0.238 (-1.04, -0.109) | 0.163 |

**Table 2.** Calculated summary statistics (median, minimum and maximum) by KRAS status in mCRC patients.

| Variable | KRAS status | Class | FOLFOX alone | Panitumumab + FOLFOX | p-value |
| --- | --- | --- | --- | --- | --- |
| Starting lesion size (mm) | Mutant | - | 38 (20, 131) | 39 (20, 150) | - |
|  | Wild-type | - | 40 (20, 190) | 38 (20, 191) | - |
| Net rate (mm/day) | Mutant | Overall | -0.111<br>(-0.504, 0.246) | -0.13<br>(-0.602, 0.323) | - |
|  |  | PD | 0.182<br>(0.132, 0.246) | 0.283<br>(0.133, 0.323) | 0.222 |
|  |  | SD | -0.104<br>(-0.292, 0.067) | -0.0835<br>(-0.219, 0.0364) | 0.263 |
|  |  | PR/CR | -0.221<br>(-0.504, -0.0848) | -0.235<br>(-0.602, -0.13) | 0.593 |
|  | Wild-type | Overall | -0.134<br>(-0.898, 0.299) | -0.18<br>(-1.04, 0.208) | - |
|  |  | PD | 0.197<br>(0.0884, 0.299) | 0.155<br>(0.102, 0.208) | 0.857 |
|  |  | SD | -0.0698<br>(-0.435, 0.241) | -0.0699<br>(-0.368, 0.0585) | 0.905 |
|  |  | PR/CR | -0.217<br>(-0.898, -0.107) | -0.252<br>(-1.04, -0.109) | 0.145 |

**Table 3.**
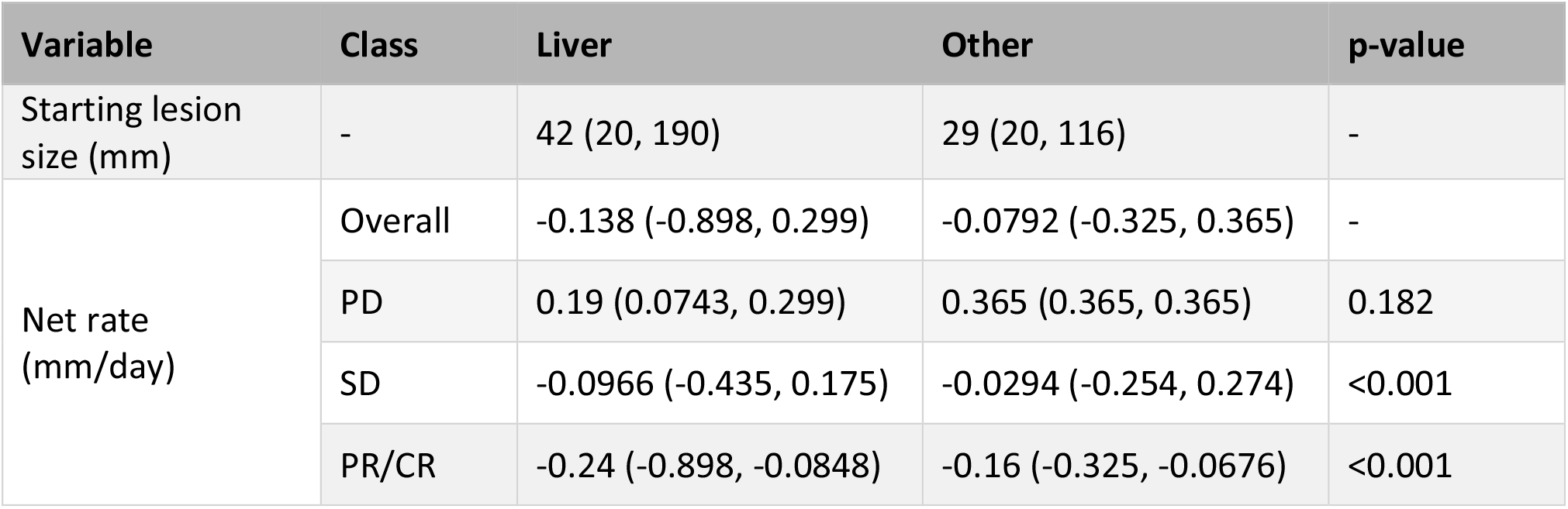
Calculated summary statistics (median, minimum and maximum) by lesion site in mCRC patients treated with FOLFOX alone.

**Table 4.** Calculated summary statistics (median, minimum and maximum) by lesion site in mCRC patients treated with Panitumumab + FOLFOX.

| Variable | Class | Liver | Other | p-value |
| --- | --- | --- | --- | --- |
| Starting lesion size (mm) | - | 41 (20, 191) | 32 (20, 110) | - |
| Net rate (mm/day) | Overall | -0.165 (-1.04, 0.395) | -0.109 (-0.54, 0.204) | - |
|  | PD | 0.283 (0.101, 0.395) | 0.0714 (0.0217, 0.0789) | 0.017 |
|  | SD | -0.0862 (-0.368, 0.0909) | -0.055 (-0.463, 0.204) | 0.174 |
|  | PR/CR | -0.259 (-1.04, -0.0923) | -0.163 (-0.54, 0.069) | <0.001 |

**Figure 1.**
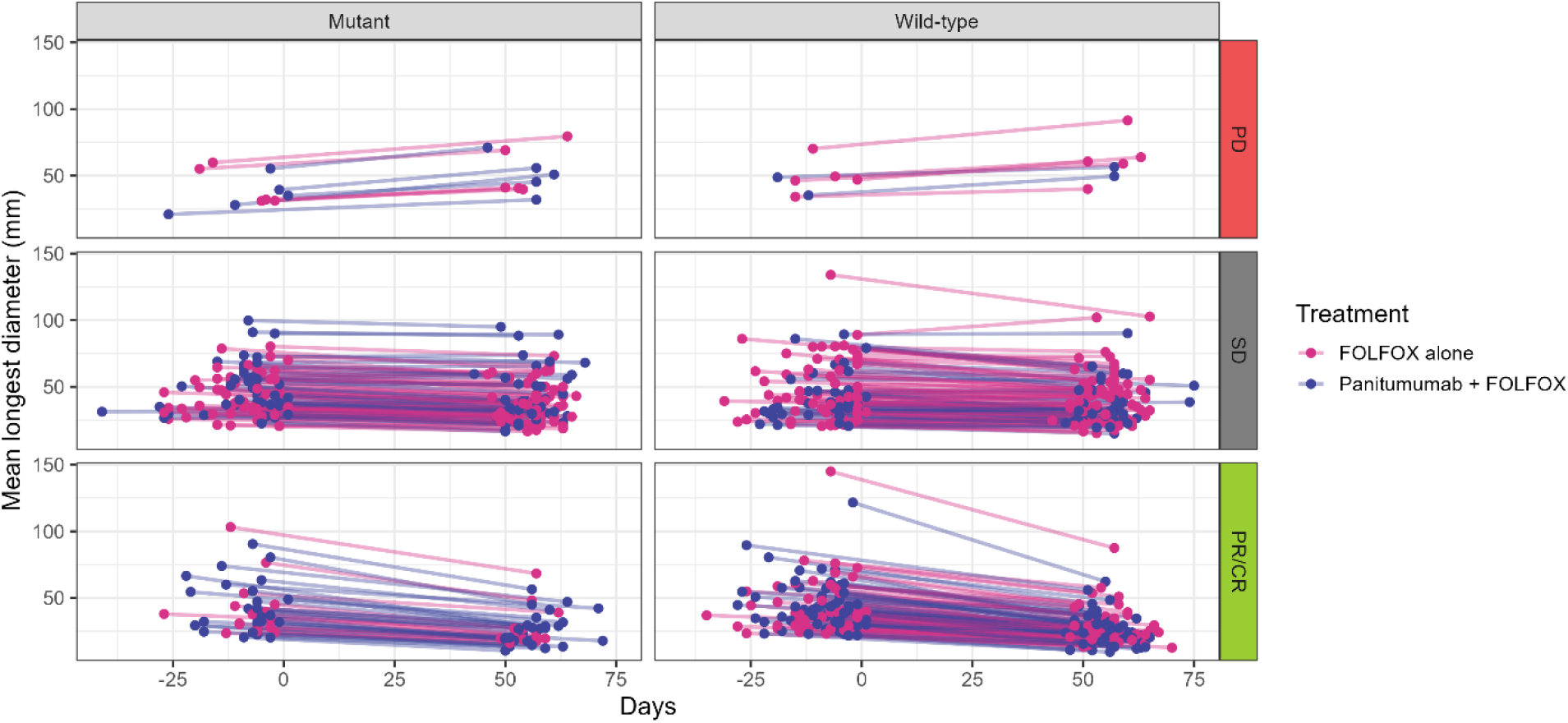
Mean longest lesion time-series data, up to the first imaging visit at Week 6, in mCRC cancer patients stratified by KRAS status.

**Figure 2.**
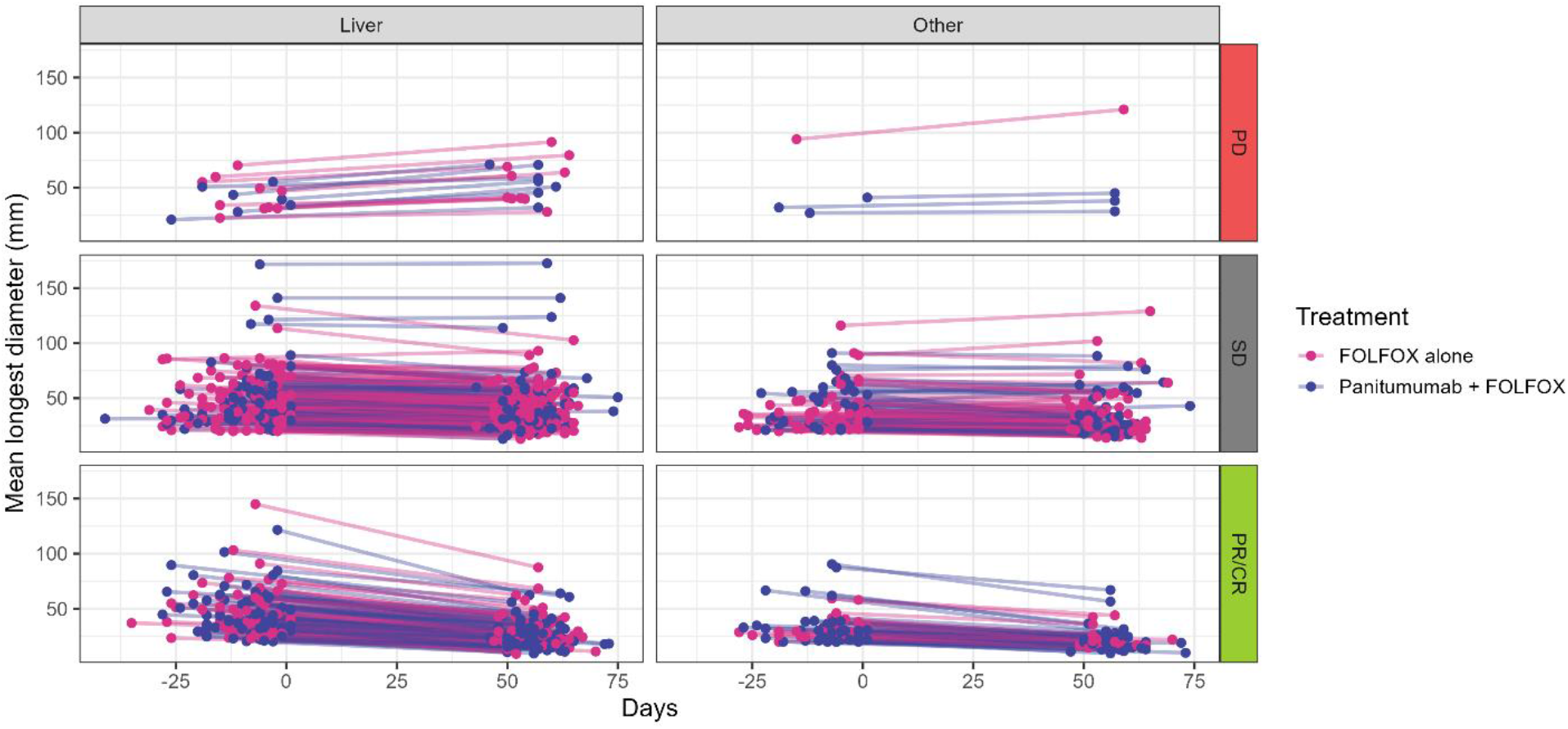
Mean longest lesion time-series data, up to the first imaging visit at Week 6, in mCRC cancer patients stratified by lesion site.

### GEJ/HCC/PC

Tables 5 to 10 present the lesion dynamics data for GEJ, HCC and PC. These tables are stratified by response classification and further stratified by liver lesions versus lesions in other locations. Similar to the study conducted on mCRC, these tables demonstrate that liver lesions that progress exhibit a faster growth rate compared to other lesions, irrespective of the primary tumour type in patients.

**Table 5.** Calculated summary statistics (median, minimum and maximum) in GEJ cancer patients.

| Variable | Class | Median (Minimum, Maximum) |
| --- | --- | --- |
| Number of lesions per patient | - | 3 (1, 12) |
| Starting lesion size (mm) | - | 22 (7, 227) |
| Net rate (mm/day) | Overall | -0.119 (-1.9, 0.833) |
|  | PD | 0.214 (-0.452, 0.833) |
|  | SD | -0.0714 (-0.595, 0.5) |
|  | PR/CR | -0.238 (-1.9, 0.19) |

**Table 6.**
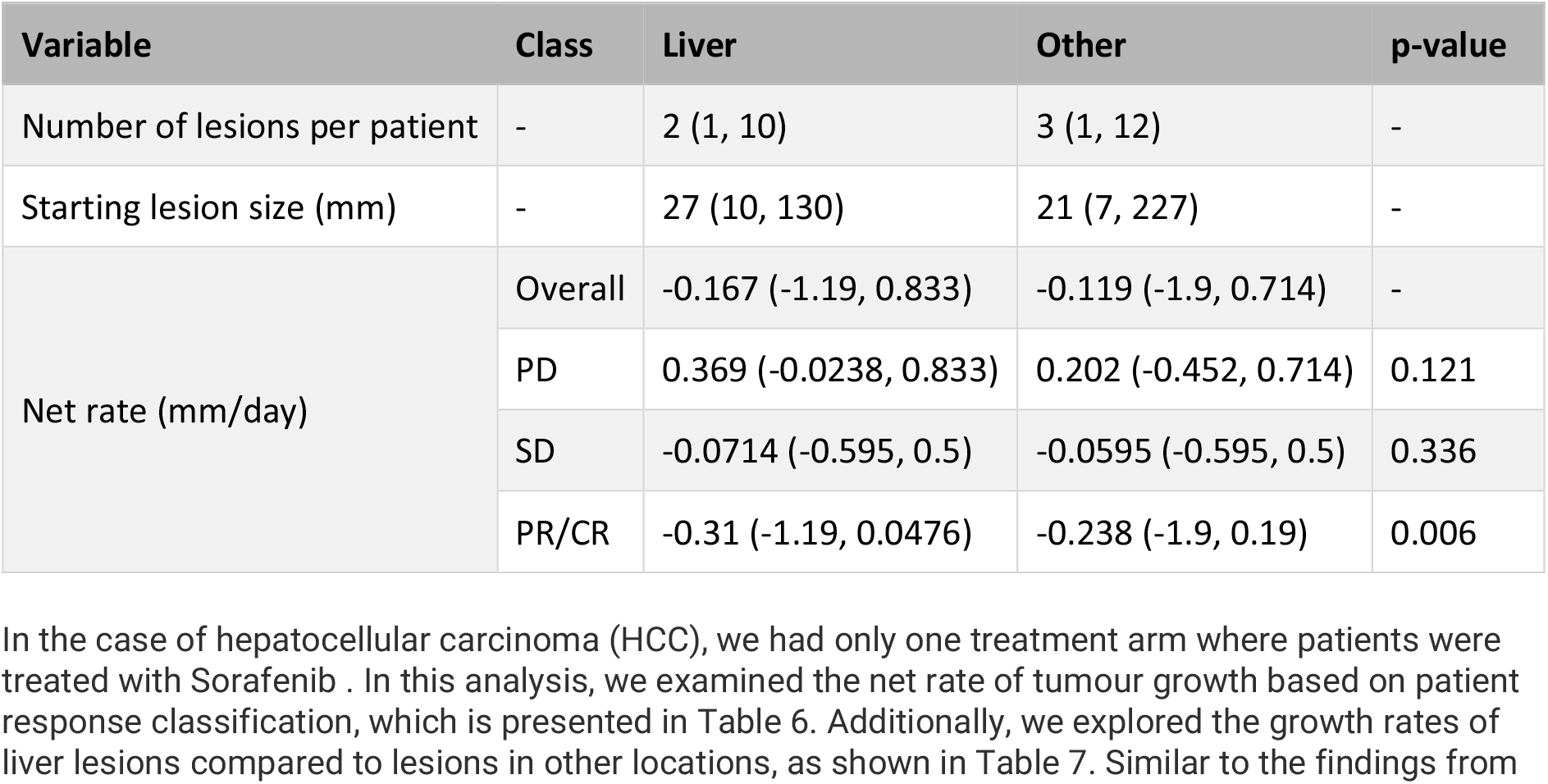
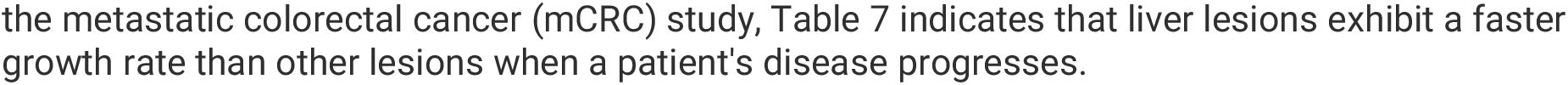
Calculated summary statistics (median, minimum and maximum) by lesion site in GEJ patients.

**Table 7.** Calculated summary statistics (median, minimum and maximum) in HCC patients.

| Variable | Class | Median (Minimum, Maximum) |
| --- | --- | --- |
| Number of lesions per patient | - | 2 (1, 10) |
| Starting lesion size (mm) | - | 27 (10, 400) |
| Net rate (mm/day) | Overall | 0 (-1.63, 9.09) |
|  | PD | 0.204 (0.0556, 9.09) |
|  | SD | 0 (-0.714, 0.87) |
|  | PR/CR | -0.2 (-1.63, -0.0755) |

**Table 8.** Calculated summary statistics (median, minimum and maximum) by lesion site in HCC patients.

| Variable | Class | Liver | Other | p-value |
| --- | --- | --- | --- | --- |
| Number of lesions per patient | - | 2 (1, 10) | 2 (1, 10) | - |
| Starting lesion size (mm) | - | 35 (10, 400) | 20 (10, 350) | - |
| Net rate (mm/day) | Overall | 0 (-1.63, 9.09) | 0 (-0.519, 0.712) | - |
|  | PD | 0.238 (0.0556, 9.09) | 0.167 (0.0566, 0.712) | <0.001 |
|  | SD | 0 (-0.714, 0.87) | 0 (-0.278, 0.224) | 0.499 |
|  | PR/CR | -0.276 (-1.63, -0.0755) | -0.163 (-0.519, -0.0769) | 0.007 |

**Table 9.** Calculated summary statistics (median, minimum and maximum) in pancreatic cancer patients.

| Variable | Class | Median (Minimum, Maximum) |
| --- | --- | --- |
| Number of lesions per patient | - | 3 (1, 8) |
| Starting lesion size (mm) | - | 24 (6, 89) |
| Net rate (mm/day) | Overall | -0.0141 (-0.436, 0.481) |
|  | PD | 0.0897 (-0.0779, 0.481) |
|  | SD | 0 (-0.219, 0.364) |
|  | PR/CR | -0.174 (-0.436, -0.109) |

**Table 10.**
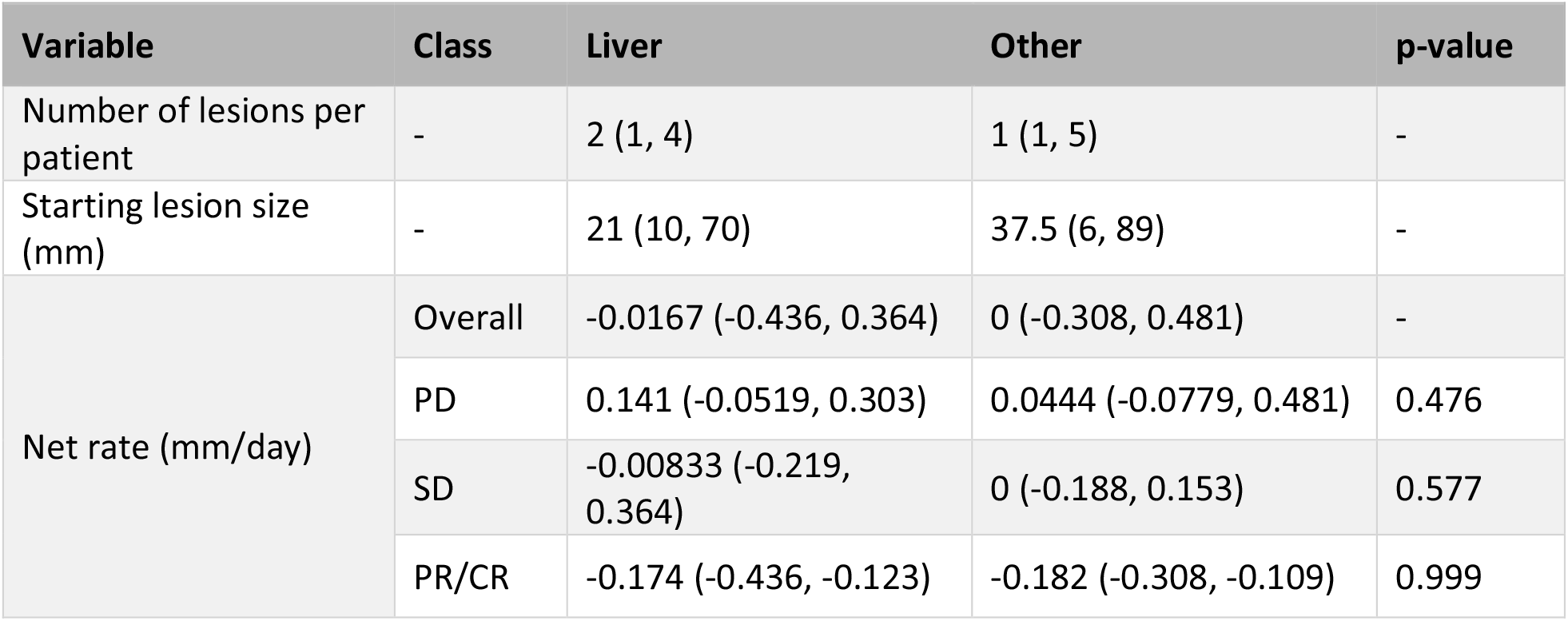
Calculated summary statistics (median, minimum and maximum) by lesion site in pancreatic cancer patients.

### NSCLC

In the studies focusing on non-small cell lung cancer (NSCLC), we have two distinct monotherapy treatments: docetaxel and erlotinib, administered within the same treatment line. Tables 11 to 14 present the tumour kinetics data, categorized first by response classification and then by liver lesions versus lesions in other locations. Consistent with our previous findings, we observe that liver lesions that progress demonstrate a more rapid growth rate compared to other progressing lesions.

**Table 11.** Calculated summary statistics (median, minimum and maximum) in NSCLC patients following treatment with Docetaxel.

| Variable | Class | Median (Minimum, Maximum) |
| --- | --- | --- |
| Number of lesions per patient | - | 3 (1, 10) |
| Starting lesion size (mm) | - | 22 (9, 180) |
| Net rate (mm/day) | Overall | 0 (-0.913, 0.882) |
|  | PD | 0.0465 (-0.55, 0.882) |
|  | SD | 0 (-0.417, 0.56) |
|  | PR/CR | -0.176 (-0.913, 0) |

**Table 12.** Calculated summary statistics (median, minimum and maximum) by lesion site in NSCLC patients following treatment with Docetaxel.

| Variable | Class | Liver | Other | p-value |
| --- | --- | --- | --- | --- |
| Number of lesions per patient | - | 1 (1, 4) | 3 (1, 10) | - |
| Starting lesion size (mm) | - | 21 (10, 77) | 22 (9, 180) | - |
| Net rate (mm/day) | Overall | 0 (-0.405, 0.553) | 0 (-0.913, 0.882) | - |
|  | PD | 0.14 (-0.184, 0.553) | 0.0366 (-0.55, 0.882) | 0.014 |
|  | SD | 0 (-0.233, 0.426) | 0 (-0.417, 0.56) | 0.775 |
|  | PR/CR | -0.148 (-0.405, -0.111) | -0.188 (-0.913, 0) | 0.721 |

**Table 13.** Calculated summary statistics (median, minimum and maximum) in NSCLC patients following treatment with Erlotinib.

| Variable | Class | Median (Minimum, Maximum) |
| --- | --- | --- |
| Number of lesions per patient | - | 3 (1, 10) |
| Starting lesion size (mm) | - | 24 (9, 190) |
| Net rate (mm/day) | Overall | 0.0141 (-0.732, 1.36) |
|  | PD | 0.179 (0.0385, 1.36) |
|  | SD | 0 (-0.554, 0.357) |
|  | PR/CR | -0.175 (-0.732, -0.0649) |

**Table 14.** Calculated summary statistics (median, minimum and maximum) by lesion site in NSCLC patients following treatment with Erlotinib.

| Variable | Class | Liver | Other | p-value |
| --- | --- | --- | --- | --- |
| Number of lesions per patient | - | 1 (1, 6) | 2 (1, 10) | - |
| Starting lesion size (mm) | - | 26 (10, 170) | 24 (9, 190) | - |
| Net rate (mm/day) | Overall | 0.0714 (-0.297, 0.957) | 0.0126 (-0.732, 1.36) | - |
|  | PD | 0.277 (0.0806, 0.957) | 0.164 (0.0385, 1.36) | <0.001 |
|  | SD | 0 (-0.297, 0.194) | 0 (-0.554, 0.357) | 0.144 |
|  | PR/CR | -0.159 (-0.273, -0.0698) | -0.175 (-0.732, -0.0649) | 0.326 |

## Discussion

The presence of liver metastases is known to be a poor prognostic factor in numerous cancer types, including NSCLC (8,14), GEJ (21) and mCRC(9). However, an assessment as to whether the dynamics of liver lesions is different to other types across major cancers has not been conducted previously. Here we sought to use open trial databases to assess if the dynamics of liver lesions are indeed different to those lesions in other locations under different treatment modalities.

RECIST imaging data at the individual lesions level from phase II/III trials for five tumour types, mCRC, GEJ, HCC, NSCLC and PC under different treatments was extracted from ProjectDataSphere. We calculated the initial net-rate of change, defined as the difference in size between pre and first post-treatment time-point divided by the time between measurements, of all individual lesions across all studies. We found evidence that lesions that are classified as progressing according to RECIST had higher growth rates if they were liver lesions versus other types across all studies bar pancreatic cancer, see Figure 3. Thus, suggesting that when treatments fail, liver lesions progress rapidly.

**Figure 3.**
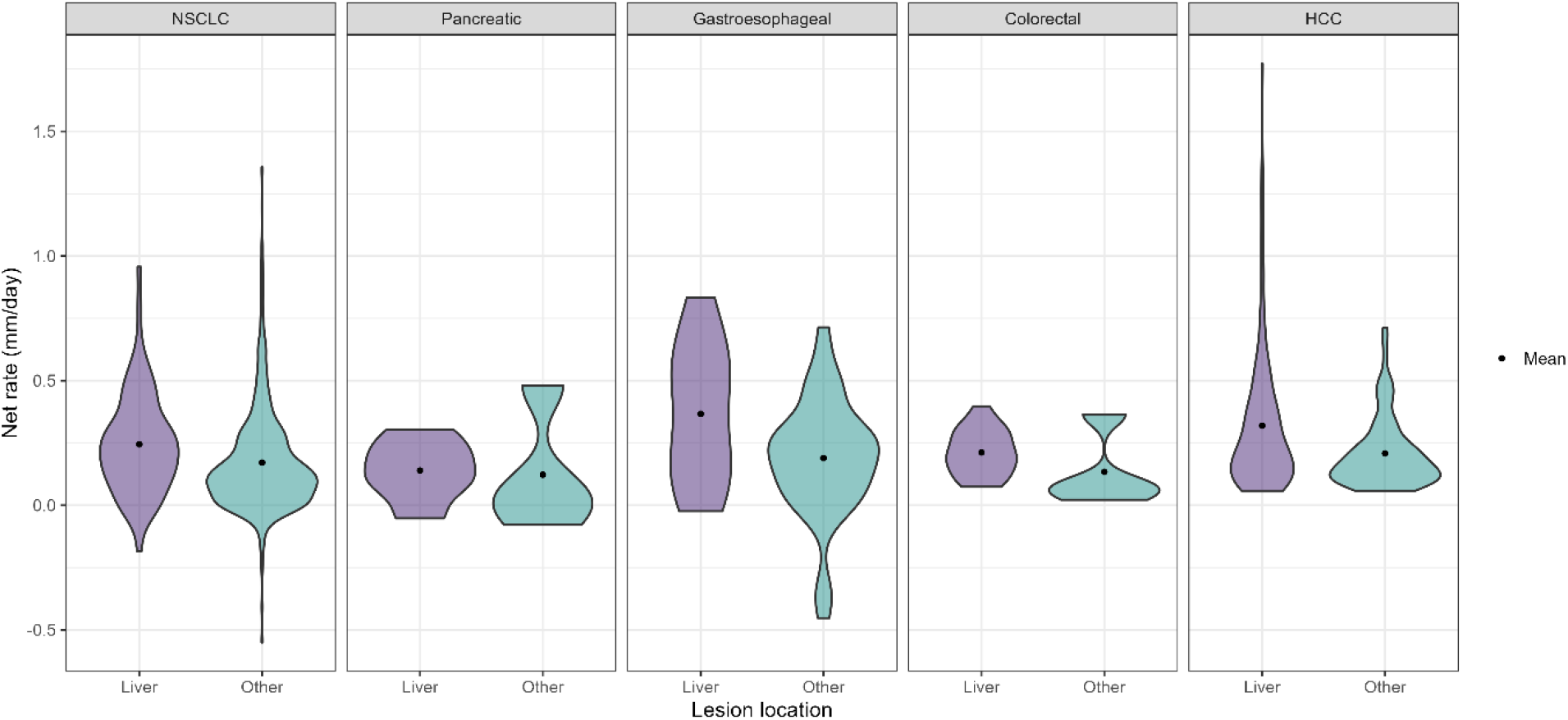
Comparison of metastases net rates across tumour sites for PD classified lesions.

These analyses are not without limitations. We do not have replicate studies to confirm our findings within the same disease/treatment type. However, given that the findings are consistent across disease/treatment modalities suggests that this may not be a huge concern. In addition, it’s known that liver metastases are a key prognostic factor thus our results confirm those inferences by highlighting these metastases grow faster than other types. Another limitation could be the quantification of the liver metastases but again our findings were consistent across numerous studies thus, if uncertainty in the measurement of lesions was an issue we would not have seen consistent results. One key limitation which cannot be addressed is that none of our studies involved immunotherapy treatments, but it must be noted that liver metastases are emerging to be a key prognostic factor for those treatments too (14).

Since liver metastases are known to be a key prognostic factor our results suggest that more basic science research is need to understand the micro-environmental niche that allows liver lesions to progress so rapidly versus lesions in other sites. The unique environment of the liver clearly provides cancer cells sufficient nutrients to grow at a faster rate than at other sites. Little work has been done on understanding the mechanisms of liver metastasis progression and specific treatments have yet to emerge to tackle this key metastatic site. Our analyses argue that more should be done and development of new treatments that target liver metastases are warranted.

Finally, our analyses may also be useful for those developing mathematical models of tumour growth and response as we have provided key summary statistics that could be used directly within such models. The analyses give distribution of both lesion size and rates which are key parameters in numerous models of tumour growth and response. These could be used to assist in back translational approaches (22) which have been used to set targets for drug development teams when selecting compounds.

In summary, the analyses presented here we hope will encourage basic scientists to explore the metastatic niche of the liver and hopefully continue to encourage organisation to deposit more clinical trial data into the open databases.

## Acknowledgements

This publication is based on research using information obtained from www.projectdatasphere.org, which is maintained by Project Data Sphere. Neither Project Data Sphere nor the owner(s) of any information from the web site have contributed to, approved or are in any way responsible for the contents of this publication.

